# Metabolomic analyses of plasma and liver of mice fed with immature *Citrus tumida* peel

**DOI:** 10.1101/430181

**Authors:** Atsushi Toyoda, Mizuho Sato, Masaki Muto, Tatsuhiko Goto, Yuji Miyaguchi, Eiichi Inoue

**Affiliations:** College of Agriculture, Ibaraki University, Ami, Ibaraki 300–0393, Japan; Ibaraki University Cooperation between Agriculture and Medical Science (IUCAM), Ami, Ibaraki 300–0393, Japan; United Graduate School of Agricultural Science, Tokyo University of Agriculture and Technology, Fuchu-city, Tokyo 183–8509, Japan

**Author notes:** Corresponding Author: (AT). Present address: Obihiro University of Agriculture and Veterinary Medicine, Obihiro, Hokkaido 080–8555, Japan (Tatsuhiko Goto).

## Abstract

Supplementing food with functional small molecules has been shown to prevent diseases and improve the quality of life, especially in elderly people. Citrus fruits and citrus fruit-products are popular food supplements across the world. In this study, we focused on a Japanese citrus fruit, *Citrus tumida* hort. ex Tanaka (*C. tumida*), and elucidated the effects of supplementation of the peels of immature *C. tumida* (PIC) on food intake, body and fat tissue weights, and metabolic profiles of plasma and liver in mice. Supplementation with 5% (w/w) PIC for 4 weeks significantly suppressed body weight gain and decreased adipose tissue weight, including that of the epididymal, perirenal, and subcutaneous fats. Metabolome analyses using capillary electrophoresis time-of-flight mass spectrometry showed that the level of 2-hydroxyvaleric acid was reduced in the blood plasma of mice fed with PIC. Supplementation with PIC significantly elevated the levels of dipeptides (Thr-Asp, Ser-Glu, and Ala-Ala), glucuronic acid (and/or galacturonic acid-2), and S-methylglutathione, and significantly reduced the levels of betaine aldehyde in the liver. In conclusion, PIC supplementation affects the metabolism of fatty acids, pectin, glutathione, and choline. Our study demonstrates the potential beneficial effects of PIC, especially in metabolic syndrome and obesity. PIC may be developed as a functional food and used in the treatment of these diseases. Nutritional and metabolome studies are effective in studying the effects of specific dietary supplements and will contribute to the development of functional foods.

## Introduction

Preventive and alternative medicines have shown to improve the quality of life, especially in the elderly population. Good health conditions may be attained by life habits, such as regular exercise, good sleep, and a proper diet. The effects of food on the quality of life have been extensively studied, and various functional foods and supplements are recommended to preserve or improve health. Fruits contain potent ingredients that affect our health. Citrus fruits are popular in several countries and have various health benefits [1]. The dried peels of *Citrus unshiu* and *C. reticulata*, which are used as natural medicines in Japan and China, show beneficial effects, including improved brain function [2]. The peel extract of *C. depressa* helps prevent obesity in mice fed with a high-fat diet [3]. Moreover, supplements of *C. unshiu* peel extract have been found to restore adenocarcinoma-induced weight loss in mice [4]. Citrus peels contain high amounts of flavonoids, which contribute to the health-beneficial effects [2]. Hesperidin and nobiletin flavonoids found in several citrus fruits show beneficial effects against some features of metabolic syndrome [1]. Interestingly, nobiletin shows a protective effect against metabolic syndrome by enhancing circadian rhythms [5], and oral administration of hesperidin reduces the levels of inflammatory markers in patients with metabolic syndrome [6]. In addition, the peel of immature citrus fruits contains relatively high levels of flavonoids and antioxidants than those in mature fruit peels [7,8].

*C. tumida* hort. ex Tanaka is a native citrus found around Mt. Tsukuba in Ibaraki prefecture, Japan [9]. *C. tumida* contains high levels of hesperidin and nobiletin than those in other species of citrus fruits [10]. However, the potential health benefits of *C. tumida* have not been investigated. Therefore, we aimed to elucidate the health benefits of *C. tumida, especially for preventing obesity and depression, which are* linked to inflammation. We first carried out a comprehensive metabolite analysis of *C. tumida*. Omics approaches are considered valuable tools to study the effects of food and farm products on health [11]. Metabolomics is frequently employed to elucidate food-induced alterations in global metabolism in humans and other animals. In this study, we investigated the health benefits of the peel of immature *C. tumida* (PIC) by evaluating the effects of diets supplemented with PIC on food intake (FI), body weight gain (BWG), and the metabolome in plasma and liver of mice.

## Materials and Methods

### Animals and plant materials

This study was approved by the Animal Care and Use Committee of Ibaraki University and conforms to the guidelines of the Ministry of Education, Culture, Sports, Science, and Technology (MEXT), Japan (Notification No. 71).

Male C57BL/6JJmsSlc (B6) mice (7-week-old) obtained from SLC Japan (Shizuoka, Japan) were housed at the animal facility of the College of Agriculture, Ibaraki University under a 12-h light-dark cycle (light on at 8:00 am). Prior to the experiments, the mice were individually housed in cages (143 × 293 × 148 mm, Charles River Laboratories Japan, Kanagawa, Japan) with wood chips. The mice were fed with a semi-purified diet (AIN-93G, Oriental Yeast, Tokyo, Japan). Food and water were available *ad libitum* and were weighed to monitor the daily consumption. Body weight was also determined daily to calculate BWG.

Immature C. tumida was harvested at an orchard in the eastern foothill of Mt. Tsukuba, Ibaraki prefecture, Japan, in early October 2015. Peels containing the outer orange layer and the inner white layer were manually collected and freeze-dried using a freeze-dryer (FDU-1110, TOKYO RIKAKIKAI, Tokyo, Japan). Dried peels were powdered using a centrifugal mill (ZM-1, Retsch technology GmbH, Haan, Germany).

Peel powder was stored at room temperature (23–26 °C) until use.

### Detection of flavonoids by high-performance liquid chromatography

To determine the flavonoid levels in PIC, concentrations of nobiletin, narirutin, geosmin, hesperidin, and tangeretin were simultaneously analyzed by high-performance liquid chromatography (HPLC) (Hitachi Chromaster System, Hitachi, Tokyo, Japan). Flavonoid standards were purchased from Wako Pure Chemical Industries, Ltd (Osaka, Japan). Dried PIC powder (100 mg) was mixed and extracted with 4 mL methanol:dimethyl sulfoxide (1:1) with agitation for 12 h at room temperature using a shaker (NR-10, Taitec, Tokyo, Japan). After the eluted solution was centrifuged at 1000 *g* for 5 min, the supernatant was collected and filtered through a membrane filter (Millex-GS 0.22 μm, Merck Millipore, Darmstadt, Germany). Samples were stored at −80 °C until analysis. A 10-μL sample aliquot was injected into the HPLC apparatus and analyzed with a photodiode array detector. ZORBAX SB-C8 (150 × 3.0 mm i.d.) (Agilent Technologies, Tokyo, Japan) was used as a separation column, and Agilent Hardware Kit High Press was used as a guard column (Agilent Technologies). The temperature of the column oven was set at 40 °C and spectra from 200 to 450 nm were obtained. The linear gradient elution program consisted of an initial 20 min (mobile phase, from 80% and 20% to 0% and 100% of formic acid and methanol, respectively) followed by 5 min of 100% methanol at a flow rate of 1.0 mL/min. Concentrations of the compounds were calculated from integrated peak areas of the sample and the corresponding authentic standards.

### Experimental design

After acclimatization to the environment of the animal facility in Ibaraki University for one week, the B6 mice were divided into two groups: control (*n*= 6) and PIC-fed (*n* = 7). The PIC-fed group was fed 5% (w/w) PIC powder in the AIN93G powder diet, whereas the control group was fed only AIN93G powder for four weeks.

### Tissue sampling

After fasting from 9:00 am to 12:00 am, the mice were sacrificed by decapitation and trunk blood was collected in a tube on ice. Final concentration was set at 0.13% EDTA-2K. The sample was centrifuged at 1,200 *g* at 23 °C for 10 min. The supernatant blood plasma was collected and stored at −80 °C until use. Approximately 50 mg from the left lobe of the liver was removed and immediately frozen in liquid nitrogen and stored at −80 °C until use. Epididymal, perirenal, and subcutaneous fats were collected and weighed. Tissue weight was normalized to body weight (BW) on the day of sampling.

### Metabolomic analysis of plasma and liver

Nine representative mice were selected for the metabolome analysis. The plasma and liver samples (*n*= 5 in the control group, *n*= 4 in PIC-fed group) were subjected to metabolomic analysis as previously reported [12]. Sample preparation and metabolome analysis were carried out by HMT (Human Metabolome Technology Inc. Tsuruoka, Japan). Capillary electrophoresis time-of-flight mass spectrometry (CE-TOFMS) analysis of the metabolome was performed using an Agilent CE Capillary Electrophoresis System equipped with an Agilent 6210 Time of Flight mass spectrometer (Agilent Technologies, Waldbronn, Germany) at HMT following previously described protocols [12–15]. Briefly, approximately 50 mg frozen liver sample was immersed in 1800 μL 50% acetonitrile in Milli-Q water (Millipore-Japan, Tokyo, Japan) containing internal standards (H3304–1002, HMT). The tissue was homogenized using the BMS-M10N21 homogenizer (BMS, Tokyo, Japan) and then centrifuged at 2300 *g* for 5 min at 4 °C. Next, 800 μL of the upper layer was filtered by centrifugation using an HMT 5-kDa cut-off filter (UFC3LCCNB-HMT, HMT) at 9100 *g* for 120 min at 4 °C. The filtrate was resuspended in 50 μL Milli-Q water for CE-MS analysis. For plasma, 50 μL sample was added to 450 μL methanol containing internal standards (H3304–1002, HMT). The solution was mixed with 500 μL chloroform and 200 μL Milli-Q water and centrifuged at 2300 *g* for 5 min at 4 °C. Next, 400 μL of the upper layer was filtered through an HMT 5-kDa cut-off filter as described above. The filtrate was then resuspended in 25 μL Milli-Q water for CE-MS analysis.

The identified metabolites from the metabolome library were assigned to the Kyoto Encyclopedia of Genes and Genomes (KEGG), facilitating the search for the corresponding metabolic pathways [16].

### Statistical analysis

BWG, FI, water intake (WI), and tissue weights were analyzed by an unpaired two-tailed Student’s *t*-test. Data were analyzed using Excel (Microsoft, WA) and are shown as mean ± SEM. For metabolomic analyses, Welch’s *t*-tests were used to compare the “supplementation” factor. To control for *P-values of multiple* comparisons, the false discovery rate was determined based on previously published studies [17,18]. The significance threshold was set to *Q*< 0.1.

## Results

### Concentrations of major flavonoids in PIC

We analyzed the major flavonoids, including nobiletin, narirutin, geosmin, hesperidin, and tangeretin, in PIC using HPLC (Table 1). Unfortunately, nobiletin and narirutin could not be separated under the HPLC conditions described above. Previous data showed that the concentration of nobiletin is approximately 3 times that of narirutin in *C. tumida* [41]; hence, nobiletin is relatively a major flavonoid in PIC.

**Table 1.** Flavonoid content in peels of immature *Citrus tumida* (100 mg).

| Flavonoids | (mg) |  |  |
| --- | --- | --- | --- |
| Nobiletin + Narirutin | 0.674 | ± | 0.015 |
| Geosmin | 0.078 | ± | 0.003 |
| Hesperidin | 0.206 | ± | 0.010 |
| Tangeretin | 0.249 | ± | 0.009 |

### BWG, FI, WI, and feed efficiency

We measured cumulative BWG, FI, and WI to evaluate the effect of PIC supplementation. Mice from the PIC-fed group showed lower BWG (control: 4.45 ± 0.47g vs. PIC: 3.20 ± 0.22 g, *P*= 0.0267, Fig 1A) and a tendency for lower FI (control: 126.18 ± 3.69 g vs. PIC: 117.76 ± 1.70 g, *P*= 0.0517, Fig 1B) when compared to control mice. No significant difference was observed in WI (control: 121.40 ± 4.52 g vs. PIC: 125.10 ± 1.90 g, *P*= 0.4470, Fig 1C). Mice from the PIC-fed group also showed a tendency to have a lower feed efficiency than that in control mice (control: 0.035 ± 0.0036 vs. PIC: 0.027 ± 0.0020, *P*= 0.0706, Fig 1D).

**Fig. 1.**
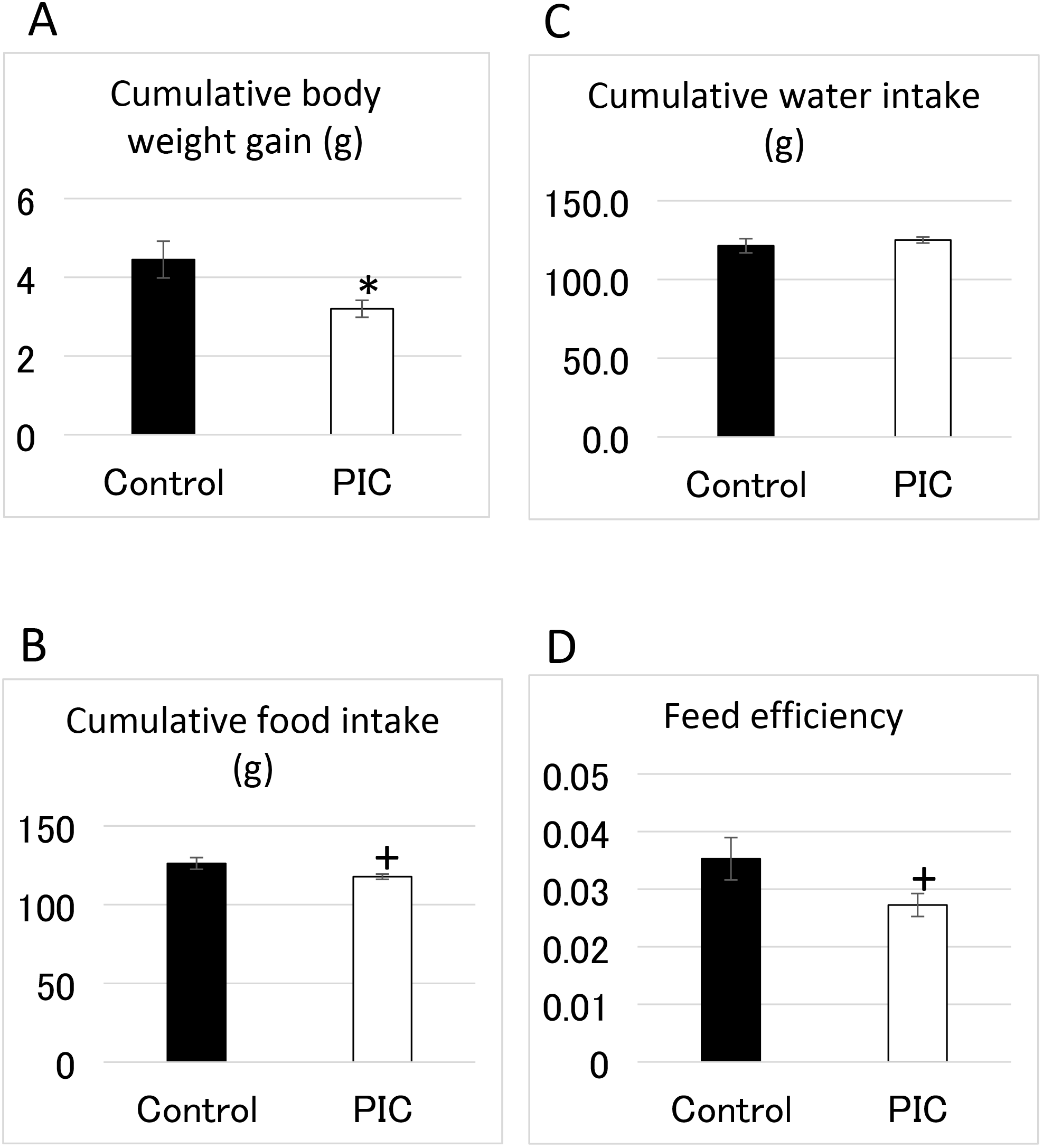
Effects of supplementation of the peels of immature *Citrus tumida* (PIC) on (A) cumulative body weight gain, (B) food intake, (C) water intake, and (D) feed efficiency (n = 6–7 in each group). Control: mice fed with AIN93G diet; PIC: mice fed with AIN93G supplemented with 5% PIC (w/w) diet. Data are expressed as mean ± SEM.^†^*P*< 0.10, *P < 0.05, \*\**P*< 0.01 versus control.

### Liver and fat weights

As shown in Fig 2A, liver weight (as % of BW) of PIC-fed mice was higher than that of control mice (control, 4.01 ± 0.13% vs. PIC, 4.57 ± 0.057 %, *P*= 0.0015). The weight of epididymal, perirenal, and subcutaneous fats (as % BW) in PIC-fed mice was significantly lower than that in control mice (epididymal fats – control: 1.85 ± 0.15 % vs. PIC: 1.40 ± 0.047 %, *P*= 0.0104; perirenal fat – control: 0.60 ± 0.067 % vs. PIC: 0.35 ± 0.010 %, *P = 0.0021; subcutaneous fat – control, 0.32 ± 0.026 % vs. PIC, 0.22 ± 0.029*%, *P = 0.0404) (Fig 2B, 2C, and 2D)*.

**Fig. 2.**
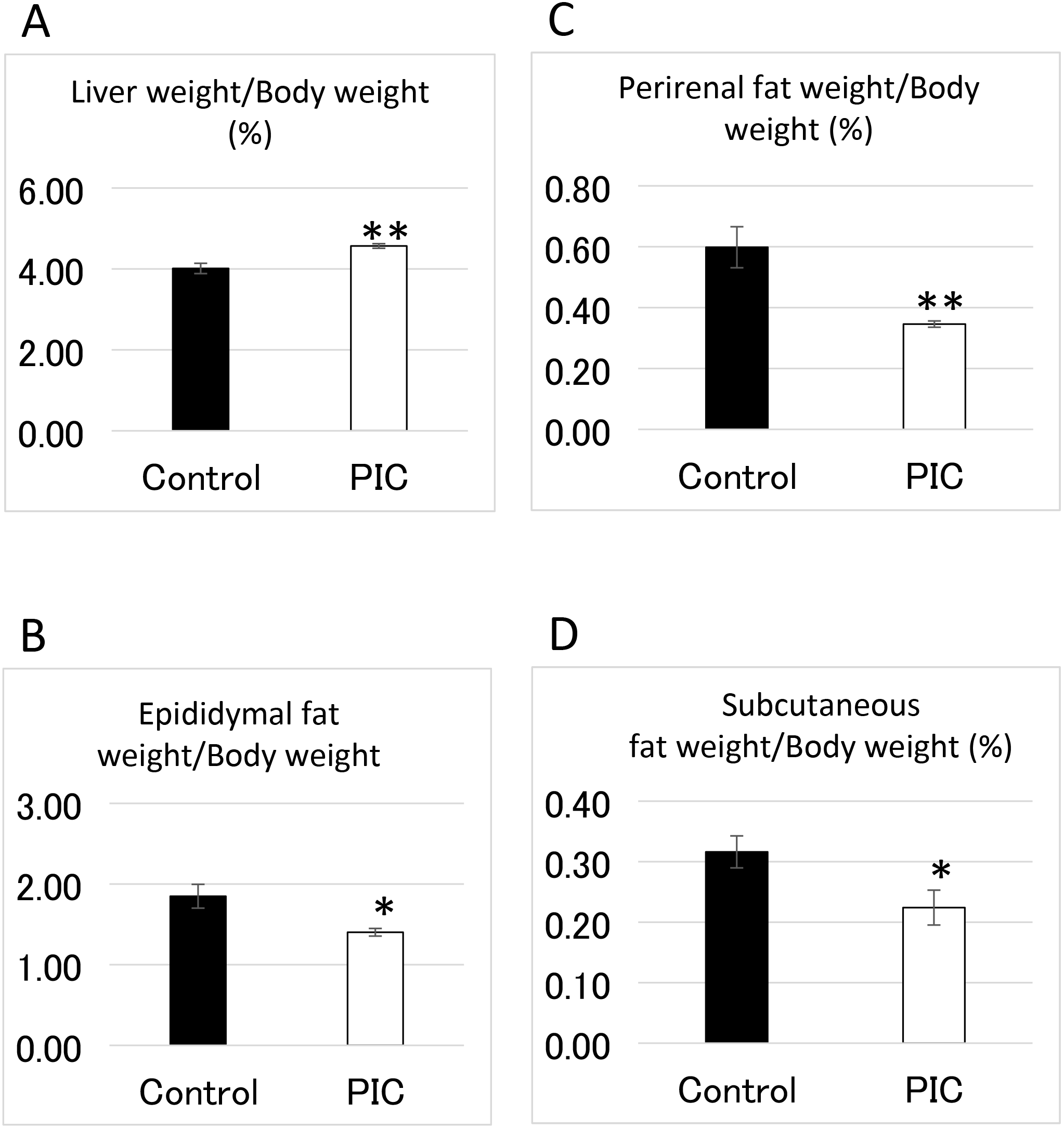
Effects of supplementation of the peels of immature *Citrus tumida* (PIC) on (A) cumulative liver weight, (B) epididymal fat weight, (C) perirenal fat weight, and (D) subcutaneous fat weight (n = 6–7 in each group). Control: mice with fed AIN93G diet; PIC: mice fed with AIN93G containing 5% PIC (w/w) diet. Data are expressed as mean ± SEM.\**P*< 0.05, \*\**P*< 0.01 versus control.

### Metabolomics

CE-TOFMS revealed 191 metabolites in the plasma. A single metabolite, 2-hydroxyvaleric acid, showed significantly (*P < 0.05 and Q*< 0.1) lower levels in PIC-fed mice than in control mice (Table 2).

**Table 2.** Metabolites affected by ingestion of peels of immature *Citrus tumida* (*Q*<0.1).

| metabolite | sample | ratio<br>(supl/cont) | *p* | *Q* |
| --- | --- | --- | --- | --- |
| 2-Hydroxyvaleric acid | plasma | 0.8 | 6.0326E-05 | 0.01152 |
| Thr-Asp<br>(Ser-Glu) | liver | 2.4 | 1.1E-04 | 0.02762 |
| Daminozide<br>(Ala-Ala) | liver | 1.4 | 4.5E-04 | 0.05596 |
| Glucuronic acid<br>(Galacturonic acid) | liver | 1.6 | 4.6E-04 | 0.03793 |
| Betaine aldehyde | liver | 0.5 | 8.3E-04 | 0.05161 |
| S-Methylglutathione | liver | 1.6 | 9.1E-04 | 0.04538 |

A total of 250 metabolites were detected by CE-TOFMS in the liver. As shown in Table 1, 5 metabolites showed significant differences in levels between PIC-fed and control mice (P < 0.05 and Q < 0.1). The relative amounts of Thr-Asp (and/or Ser-Glu), daminozide (and/or Ala-Ala), glucuronic acid (and/or galacturonic acid-2), and S-methylglutathione were significantly higher in PIC-fed mice than in control mice. In contrast, the relative amount of betaine aldehyde was significantly lower in PIC-fed mice than in control mice.

## Discussion

In this study, we investigated the health benefits of a local citrus fruit, *C. tumida*, and focused on the dietary effects of PIC, which contains flavonoids, such as nobiletin and hesperidin. As shown in Table 1, PIC contained high amounts of nobiletin and narirutin compared to other flavonoids, although the peaks of nobiletin and narirutin could not be separated under the conditions described in the Methods. Supplementation of PIC significantly suppressed BWG compared to that observed with the control diet (*p* < 0.05, Fig 1A); although FI and feed efficiency were reduced slightly, the difference was not statistically significant (*p* < 0.1, Fig 1B and 1D). In addition, PIC intake significantly decreased the weight of adipose tissues (p < 0.05, Fig 2B, 2C, and 2D). These effects may result from the presence of flavonoids in PIC, which can affect lipid metabolism. In particular, *C. tumida* contains high levels of hesperidin, nobiletin, and tangeretin [10]. Hesperidin has several biological and pharmacological properties, such as anti-inflammatory, anti-carcinogenic, anti-oxidative, vascular protective, and lipid-lowering activities [19–21]. Moreover, nobiletin and hesperidin repress the expression of genes related to lipid synthesis, such as stearoyl-CoA desaturase [22]. In general, several citrus fruit-flavonoids have anti-inflammatory, insulin-sensitizing, and lipid-lowering activities [1]. Therefore, the flavonoids in PIC may reduce body fat deposition and suppress BWG in mice.

Metabolomics is widely used in food science and nutrition research [23], and the use of CE-TOFMS reveals the global effects of food on metabolism [11]. In this study, the plasma and liver metabolites of PIC-fed and control mice were analyzed and compared comprehensively using CE-TOFMS, and significant differences were detected, suggesting that this approach may provide important metabolic information about the effects of oral PIC supplementation. In humans, the metabolomic analysis of the effects of diets, including citrus fruits, was similarly successful [24].

The blood plasma levels of 2-hydroxyvaleric acid (2-hydroxypentanoic acid) decreased in PIC-fed mice (*p* < 0.05, *Q* < 0.1, Table 2). 2-Hydroxyvaleric acid is present in human fluids and is related to the pathophysiological metabolites in acidosis [25]. The biological function of 2-hydroxyvaleric acid in animal tissues is unclear. Non-obese diabetic (NOD) mice are widely used in type 1 diabetes studies, and they are divided into progressor and non-progressor groups depending on whether or not the animal shows disease progression [26]. The authors showed that plasma levels of 2-hydroxyvaleric acid are lower in non-progressor than in progressor NOD mice, and therefore, 2-hydroxyvaleric acid may be a good predictor of type 1 diabetes. In addition, plasma levels of 2-hydroxyvaleric acid in humans are decreased by simvastatin, a statin that reduces LDL-cholesterol levels and the risk of cardiovascular disease [27]. It is possible that simvastatin, or a metabolite of the drug, inhibits an enzyme that produces 2-hydroxyvaleric acid. Although the precise mechanism underlying these effects is unknown, PIC may modify the fatty acid metabolism and downregulate the 2-hydroxyvaleric acid synthesis pathway, resulting in lower levels in the plasma.

Thr-Asp and/or Ser-Glu levels increased in the liver of PIC-fed mice compared to those in the liver of control mice (*p* < 0.05, *Q* < 0.1, Table 2). PIC may increase proteolysis in the liver, leading to increased dipeptide levels. Daminozide and/or Ala-Ala levels also increased in the liver of PIC-fed mice (*p* < 0.05, *Q* < 0.1, Table 2). Daminozide is used as a pesticide and is unlikely to be normally present in animal tissues. Ala-Ala is a dipeptide derived from proteolysis, and increased levels may be the result of increased protein catabolism. D-Ala-D-Ala is an important component of peptidoglycan in the bacterial cell wall [28,29]. A previous study reported that vancomycin-sensitive gram-positive gut bacteria may promote hepatocellular carcinoma through the enterohepatic circulation of gut bacterial metabolites or toxins [30]. D-Ala-D-Ala in the gut may, therefore, be carried to the liver via the enterohepatic circulation. Dietary plant fibers, including pectin, a major component in citrus peels, strongly affect the intestinal bacterial ecosystem [31] and thus increase the Ala-Ala levels in the liver of PIC-fed mice.

The levels of glucuronic and UDP-glucuronic acids increased in the liver after PIC supplementation (*p* < 0.05, Table 2, Table S2). Since both of them play a pivotal role in the elimination of toxic substances [32], increased levels of glucuronic and UDP-glucuronic acids by PIC may contribute to enhanced detoxification in the liver.

PIC also increased the levels of galacturonic acid in the liver (*p* < 0.05, *Q* < 0.1, Table 2). Galacturonic acid is the main component of pectin [33], which is digested by several fibrolytic enzymes produced by intestinal microorganisms [34]. Pectin is a major component of citrus peels. Therefore, the digestion of PIC produces galacturonic acid in the mouse intestine, and this metabolite is possibly transported to and accumulated in the liver. It has been reported that orally fed pectin and galacturonic acid inhibit hepatic lipogenesis in rats [35]. Lipid metabolism should be studied in PIC-fed mice to clarify these and other issues. The functions of other dietary fibers in PIC should also be investigated.

Reduced glutathione (GSH), a major protectant against oxidative stress, is methylated in the SH group of its cysteine moiety and converted to S-methylglutathione. Levels of S-methylglutathione increased in PIC-fed mice compared to those in control mice (*p* < 0.05, *Q* < 0.1, Table 2). In addition, oxidized glutathione (GSSG) levels in the liver of PIC-fed mice were slightly higher than those in control mice (*p* < 0.1, Table S4). PIC may thus be able to increase anti-oxidative stress activity via glutathione metabolism in the liver. In fact, chronic administration of *C. unshiu* extracts reduces oxidative stress in the liver of diabetic rats by increasing glutathione levels [36]. The function of S-methylglutathione in the liver is unclear. However, in the central nervous system, S-methylglutathione is released upon hypoxia and depolarization [37] and central administration of S-methylglutathione mitigates acute stress in neonatal chicks [38]. Therefore, S-methylglutathione may play an important role in the central nervous system. Future studies are necessary to assess the function of S-methylglutathione in the liver and the effects of increased S-methylglutathione levels induced by PIC on the liver function.

Supplementation of PIC downregulated choline metabolism in the liver (Table 2, Table S2, Fig 3). GPCho is a product of the breakdown of phosphatidylcholine and is converted to choline by GPCho phosphodiesterase. Choline dehydrogenase produces betaine aldehyde from choline [39], and glycine betaine is converted from betaine aldehyde by aldehyde dehydrogenase [40]. The mechanism underlying the downregulation of choline metabolism in the liver by PIC supplementation remains unclear. Future studies will focus on evaluating the expression of genes related to choline metabolism in the liver of PIC-fed mice, and critical ingredients in PIC downregulating choline metabolism should be discovered.

**Fig. 3.**
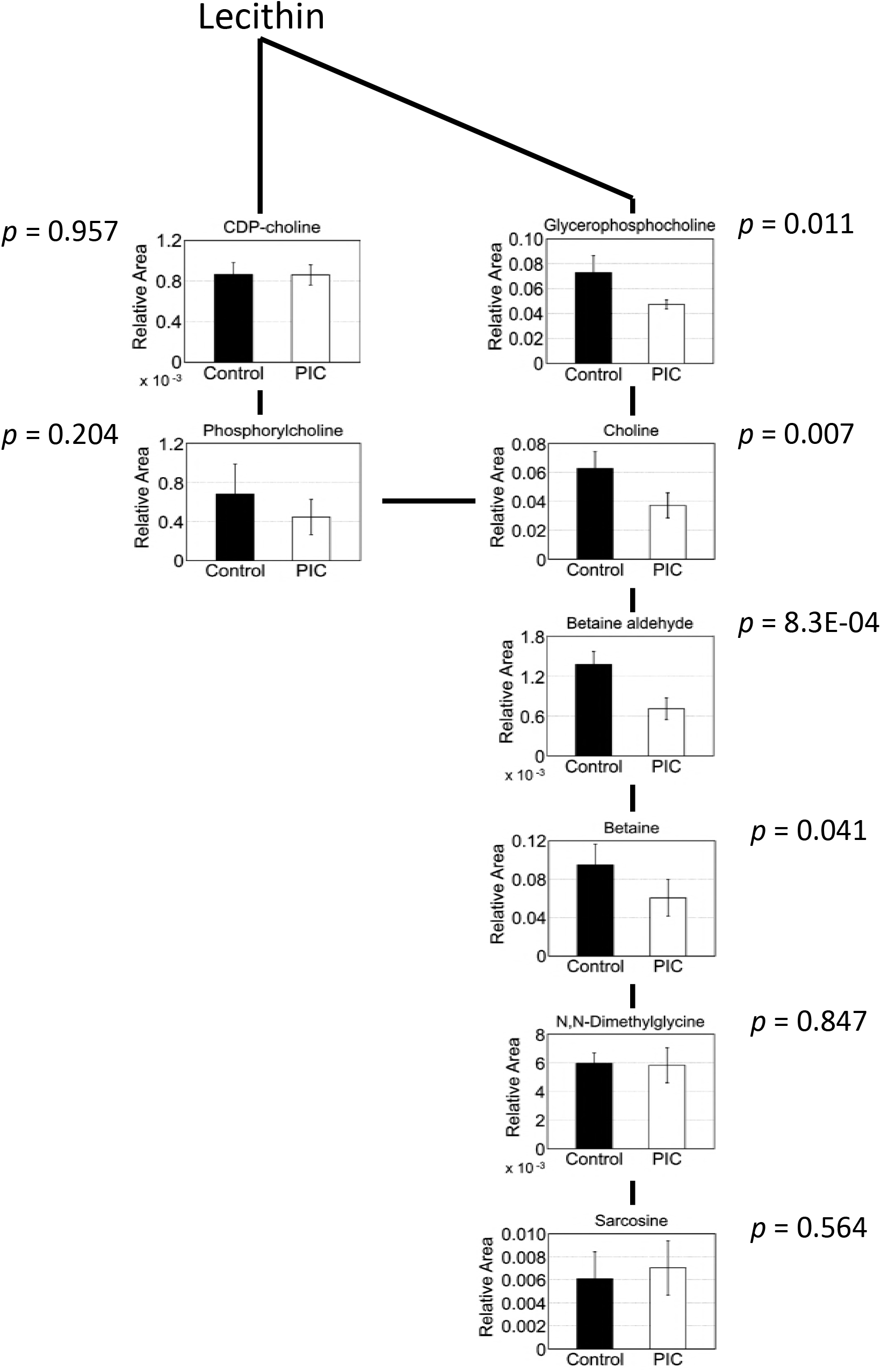
Choline-related metabolites in the liver. The pathway of choline-related metabolites in the liver is shown. Welch’s *t*-tests were used to compare “supplement,” using both peels of immature *Citrus tumida* (PIC)-fed mice (*n*= 4) and control mice (n = 5). Control: mice fed with AIN93G diet; PIC: mice fed with AIN93G containing 5% PIC (w/w) diet. Data are expressed as mean ± SEM.

In summary, we evaluated the effects of PIC supplementation on BWG, FI, WI, adipose tissue weight, and metabolomics of blood plasma and liver in mice and concluded that PIC influences body and adipose tissue weights and promotes metabolic alterations. Especially, plasma 2-hydroxyvaleric acid and liver choline metabolism including betaine aldehyde were significantly decreased by dietary PIC. In the future, we will focus on elucidating the mechanisms by which PIC supplementation reduces body fat and changes the levels of some metabolites in mice. This study used a small cohort and might have missed some critical metabolites that are significantly altered by dietary PIC. Therefore, in future studies we will try to carry out metabolomic analyses using larger cohorts of mice and humans to elucidate PIC functions, especially in fat metabolism. These studies will contribute to establish the effects of PIC on human and animal health and provide a better understanding of the health benefits of citrus fruits.

## Acknowledgments

We thank H. Shimonishi and T. Fujii (Ibaraki University) for valuable technical assistance.

## Authors’ contribution

A. Toyoda, M. Sato, and T. Goto designed the studies. A. Toyoda, M. Sato, M. Muto, Y. Miyaguchi, and E. Inoue performed the experiments. A. Toyoda and T. Goto analyzed the data. A. Toyoda wrote the manuscript.

